# Highly selective SGLT2 inhibitors suppress glucose uptake in alpha-TC1 cells, while glucagon secretion is not affected

**DOI:** 10.1101/2025.07.30.667793

**Authors:** Licht Miyamoto, Suguru Nakayama, Honoka Endoh, Kano Fujiwara, Mana Hattori, Takashi Yasuoka, Masaki Imanishi, Yasumasa Ikeda, Koichiro Tsuchiya

**Author notes:** Address correspondence and reprint requests to Licht Miyamoto, 1030, Shimo-Ogino, Atsugi-shi, Kanagawa 243-0292, Japan., (L.M.). CRedit S.N., H.E., and L.M. mainly performed experiments, and prepared figures. L.M. conceived, designed, and conducted the study, and wrote the manuscript. S.N. and K.T. also contributed to the study design. S.N., H.E. M.H., K.F., K.T., and L.M. contributed to the establishment of the assays. T.Y, M.I., and Y.I. contributed to the experiments and/or discussion. Full-length research articles. **Declaration of generative AI and AI-assisted technologies in the writing process.** Grammatical checking and rephrasing for improved readability were performed with the assistance of Perplexity.ai.

## Abstract

The existence of sodium-glucose cotransporter 2 (SGLT2) in pancreatic alpha cells and its potential roles in glucagon secretion remain controversial, despite its well-established function in renal glucose reabsorption. While some studies suggest SGLT2 presence and its involvement in glucagon regulation, others deny its expression in alpha cells.

To clarify this dispute, we investigated the functional effects of highly selective SGLT2 inhibitors, dapagliflozin and empagliflozin, on glucose uptake, intracellular ATP levels, and glucagon secretion in alpha-TC1 cells, a widely used model for glucagon-secreting cells in culture.

The SGLT2 inhibitors significantly suppressed basal glucose uptake in alpha-TC1 cells, indicating functional SGLT2 presence. However, the inhibitors did not affect glucagon secretion. Neither the SGLT2 inhibitors nor the more potent glucose transport inhibitor, cytochalasin B, altered intracellular ATP levels or glucagon secretion. In contrast, pharmacological inhibition of K/ATP channels increased glucagon secretion without affecting glucose uptake or ATP levels.

These results suggest that while SGLT2 is functionally present at low levels and mediates basal glucose uptake in alpha-TC1 cells, its inhibition has insufficient influence on intracellular ATP levels, and therefore glucagon secretion remains stable. Furthermore, our observations support predominant involvement of K/ATP channels in regulating glucagon secretion. Further studies in human purified pancreatic alpha cells in addition to islets are warranted to fully elucidate SGLT2’s role in alpha-cell physiology.

## Introduction

Sodium-glucose cotransporters (SGLTs), which belong to the solute carrier family transporters (SLC) 5A, are membrane proteins responsible for facilitating the transport of glucose and sodium ions across cell membranes, playing a pivotal role in maintaining glucose homeostasis. SGLTs utilize the electrochemical gradient of sodium ions to drive the active transport of glucose against its concentration gradient into cells. Among the more than 10 members of the SLC5A family, SGLT1 and SGLT2 stand as the most prominent and extensively studied members of the SGLT family. SGLT1, predominantly found in the small intestine, plays a critical role in the process of dietary glucose absorption. By mediating the active transport of glucose from the intestinal lumen into the epithelial cells lining the intestine, SGLT1 facilitates the efficient absorption of glucose from digested food into the bloodstream. SGLT1 is also found in other tissues, such as the kidney and heart. SGLT2, another essential member of the SLC5A family, primarily resides in the kidneys. Its primary role involves the reabsorption of glucose from the kidney filtrate back into the bloodstream, a mechanism crucial for preventing excessive loss of glucose through urine. Both SGLT1 and SGLT2 play indispensable roles in orchestrating glucose regulation within the body, albeit in different contexts. SGLT1 assumes the primary responsibility for dietary glucose absorption within the intestine, while the role of SGLT2 in renal glucose reabsorption is of paramount importance. Understanding the intricate functions of SGLT1 and SGLT2 has profound implications for glucose metabolism and its regulation, driving the development of novel therapeutic strategies for conditions like diabetes[1].

Early studies showed that pharmacological inhibition of SGLTs by phlorizin led to a significant reduction in blood glucose levels with enhanced excretion of urinary glucose. Guided by the reports on SGLT2 as a potential therapeutic target, a series of SGLT2 inhibitors have been developed for the treatment of diabetes. As expected, SGLT2 inhibitors represent a novel class of antidiabetic agents that target renal glucose reabsorption, thereby offering a distinct mechanism of action, especially independence of insulin. While this approach has been successful to varying extents, clinical experiences of ketoacidosis and rise in endogenous glucose production which could be accompanied by hyperglucagonemia have been reported [2–4]. The insulin-independent suppression of blood glucose levels seems naturally associated with glucogenic counter responses, however, possible involvement of direct actions of SGLT2 inhibitors has been proposed. Bonner et al. demonstrated that SGLT2 expression in human pancreatic alpha cells, and its inhibition by dapagliflozin promoted glucagon secretion possibly via an ATP-sensitive potassium (K/ATP) channel in human islet *ex vivo* [5]. Pedersen et al. also showed dapagliflozin-induced increase in glucagon secretion using human and murine islets, which occurred only under high glucose conditions [6]. They proposed the possible involvement of the K/ATP channel using mathematical model, supporting Bonner’s findings.

However, the concept of the direct action of SGLT2 on alpha-cell activity has been challenged by later inconsistent observations. Solini et al. failed to detect expression of SGLT2 by quantitative PCR in murine glucagonoma alpha-TC1 cells [7]. Kuhre et al. reported no expression of SGLT2 in pancreatic alpha cells through intensive investigations using human and murine islets [8]. Although the Bonner’s group lately demonstrated high heterogeneity in expression of SGLT2 in human pancreas [9], other groups also reported that SGLT2 was not present in human and murine pancreatic alpha cells [10,11]. Instead, the expression of SGLT1 in pancreatic alpha cells has been consistently confirmed, with some reports suggesting its possible involvement in glucagon secretion [7,8,10,11].

Despite the controversial roles and expression of SGLT2 in alpha cells, functional studies using isolated alpha cells have been poorly reported. Therefore, we conducted a functional evaluation of SGLT2 inhibitors on glucose uptake using alpha-TC1 cells, as it is virtually the only and widely analyzed cultured cell line model of glucagon-secreting pancreatic alpha cells isolated from murine glucagonoma [12], to elucidate the relevance of SGLT2 in pancreatic alpha cells. In this study, we investigated the effects of highly selective SGLT2 inhibitors on glucose uptake and glucagon secretion in alpha-TC1 cells.

## Materials and Methods

### Reagents

All reagents were of analytical grade and were obtained from Wako Pure Chemical (Osaka, Japan), Tokyo Chemical Industry (Tokyo, Japan), or Kanto Chemical Industry (Tokyo, Japan) unless otherwise stated. Dapagliflozin (final concentration; 1 μM) was purchased from AdooQ Bioscience (Irvine, CA), empagliflozin (final concentration; 10 μM) was obtained from Cayman Chemical (Ann Arbor, MI, US), cytochalasin B (final concentration; 100 nM) was sourced from Wako Pure Chemical, nateglinide (final concentration; 10 μM) was acquired from Namiki Shoji (Tokyo, Japan), 2-deoxy-D-glucose was obtained from Tokyo Chemical Industry, and [^3^H]-labeled 2-deoxy-D-glucose was from American Radiolabeled Chemicals Inc. (Saint Louis, MO, US).

### Cell cultures

Murine glucagonoma alpha-TC1 cells (clone 6) were obtained from the American Type Culture Collection (Rockville, MD, USA) by courtesy of Professor T. Kitamura. The cells were cultured in Dulbecco’s Modified Eagle medium containing 2 g/L glucose, 15 mM Hepes, 10% fetal bovine serum, and 1% nonessential amino acids, in a humidified atmosphere with 5% CO_2_.

### Glucose uptake activity

Glucose uptake activity was quantified by liquid scintillation counting of partially tritium-labelled 2-deoxy-D-glucose which was taken up by the cells, following the methodology outlined in our previous reports [13,14]. In summary, cells cultured in a 24-well plate were preincubated with Kreb’s Ringer bicarbonate (KRB) buffer containing 0.2% bovine serum albumin and 25 mM Hepes for 30 min. Subsequently, the cells were exposed to the specified compounds for 10 min, and then incubated for an additional 10 min in the presence of 100 μM 2-deoxy-D-glucose, including 0.1 μCi/ml tritium-labelled tracer. The 2-deoxy-D-glucose that was taken up by the cells was measured by quantifying the scintillation light emitted from the cell lysates (ALOKA LSC-7400, Nippon RayTech, Tokyo, Japan).

### Glucagon secretion

Glucagon secretion was assessed by measuring the levels of released glucagon in the culture media during a 2-hour incubation period. Initially, the cells were preincubated in KRB buffer containing 0.2% bovine serum albumin, 25 mM Hepes, and 16.7 mM glucose for 1 hour. Subsequently, the cells were exposed to the specified compound in KRB buffer with 0.2% bovine serum albumin, 25 mM Hepes, and 5 mM glucose for 2 hours. The glucagon concentration in the buffer was determined using a glucagon assay kit that utilizes the homogenous time-resolved fluorescence (HTRF) technology (CIS Bio International, Saclay, France).

This method is based on the fluorescence resonance energy transfer (FRET), and employs two distinct antibodies targeting the N- and C-terminus of murine glucagon. Afterward, the remaining cells were lysed, and the protein content in the lysate was quantified.

### Intracellular ATP levels

ATP levels were determined as described previously[15]. The ATP levels within the cells cultured in a 96-well plate were quantified using the D-luciferin-luciferase luminescence method with 1-step ATPlite reagent (PerkinElmer, Inc., Waltham, MA) and measured using a multimode plate reader, EnSpire (Perkin Elmer, Shelton, CT). This measurement was performed following the 30-min exposure to the indicated compounds in the KRB buffer containing 0.2% bovine serum albumin, 25 mM Hepes, and 5 mM glucose.

### Statistical analyses

Comparisons between two groups were assessed using the unpaired Student’s t-test. When comparing multiple groups, a parametric Tukey-Kramer’s test was employed. The results are presented as means ± standard error. P-values of less than 0.05 were considered statistically significant.

## Results

### Highly selective SGLT2 inhibitors significantly inhibit glucose uptake activity in alpha-TC1 cells

We quantified the glucose uptake activity to determine the pharmacological influence of dapagliflozin and empagliflozin, highly specific SGLT2 inhibitors, on alpha-TC1 cells. At moderate dosages (1 μM and 10 μM, respectively), these inhibitors led to a significant reduction in glucose uptake activity by 10-25% (Fig. 1). These results indicate the functional presence of glucose transport through SGLT2 in the alpha-TC1 cells.

**Fig.1.**
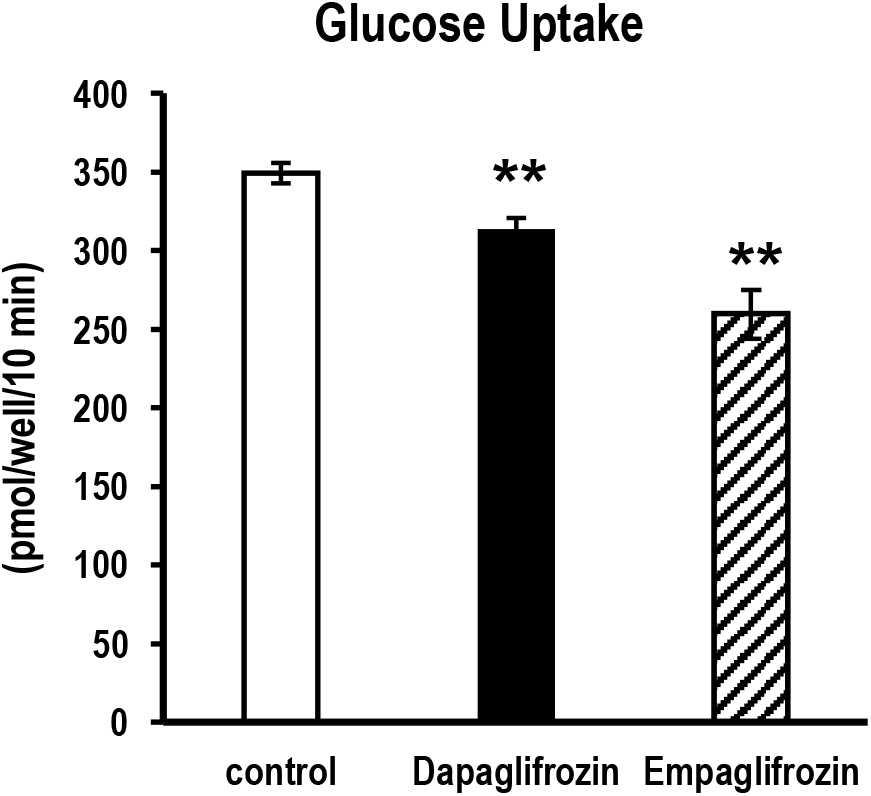
Highly selective SGLT2 inhibitors significantly suppress glucose uptake activity in alpha-TC1 cells. Uptake of 2-deoxy-D-glucose in the absence (open bar), and presence of SGLT2 inhibitors, dapagliflozin (closed bar) or empagliflozin (hatched bar). The vertical axis of the data represents the values of taken-up 2-deoxy-D-glucose per well per 10 minutes. **; p<0.01, vs control. n=4-8.

### The SGLT2 inhibitors do not affect secretion of glucagon, despite inhibiting glucose uptake

To further investigate the potential influence of SGLT2, we assessed the impact of SGLT2 inhibitors on glucagon secretion in alpha-TC1 cells. The amount of glucagon released from the cells remained unaffected by the pharmacological inhibition of SGLT2 using dapagliflozin or empagliflozin (Fig. 2).

**Fig. 2.**
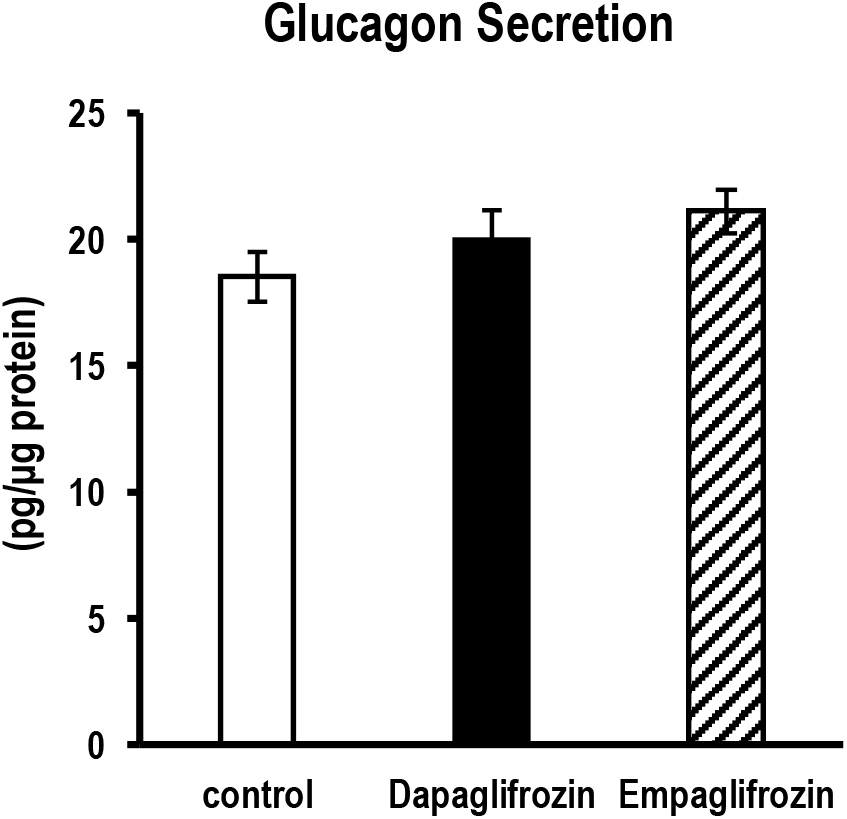
SGLT2 inhibitors do not alter glucagon secretion in alpha-TC1 cells. Glucagon secreted in the cultured media in the absence (open bar), and presence of SGLT2 inhibitors, dapagliflozin (closed bar) or empagliflozin (hatched bar). The vertical axis of the data represents the values of secreted glucagon normalized by the protein content from the cells. No significant differences vs control. n=8.

### Pharmacological inhibition of SLC2A family glucose transporters suppresses glucose uptake more intensely but does not influence glucagon secretion

The cellular glucose transport landscape comprises two distinct families of transporters. The SLC2A family, encompassing GLUT1 among others, is widely expressed in various tissues and primarily mediates basal glucose uptake. To investigate the differential impacts on glucose uptake and glucagon secretion, we employed the nonspecific inhibitor to target SLC2A family glucose transporters, cytochalasin B. The cytochalasin B exhibited a potent 60% reduction in glucose uptake activity (Fig. 3A). This robust inhibition did not lead to alterations in the amount of glucagon released from the cells (Fig. 3B).

**Fig. 3.**
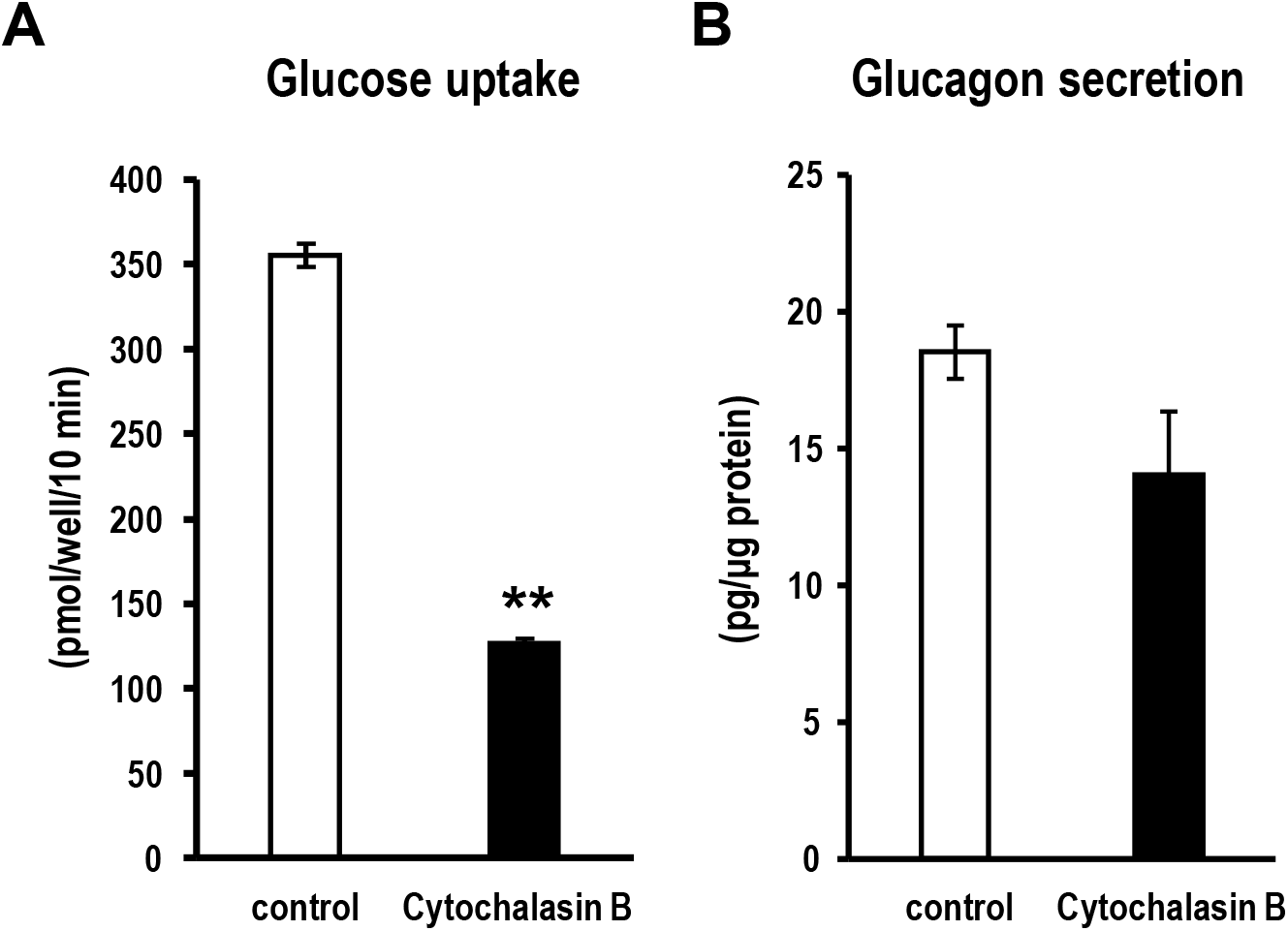
A nonspecific inhibitor of SLC2A family glucose transporters strikingly suppresses glucose uptake activity in alpha-TC1 cells, while glucagon secretion is not affected. Uptake of 2-deoxy-D-glucose (A) and glucagon secreted in the cultured media (B), in the absence (open bar) or presence (closed bar) of cytochalasin B. **; p<0.01, vs control. n=4-8.

### Pharmacological blockade of ATP-sensitive potassium channel increases glucagon secretion without impacting glucose uptake

Electrochemical excitation stands as a typical mechanism converting extracellular signals into secretion in endocrinological tissues, including pancreatic beta cells. In light of this, we examined the potential involvement of the ATP-sensitive potassium (K/ATP) channel in glucagon secretion. The pharmacological blockage of K/ATP channel using a weak agonist targeting the sulfonylurea receptor led to an increase in glucagon secretion, while glucose uptake activity remained unchanged (Fig. 4A, B).

**Fig. 4.**
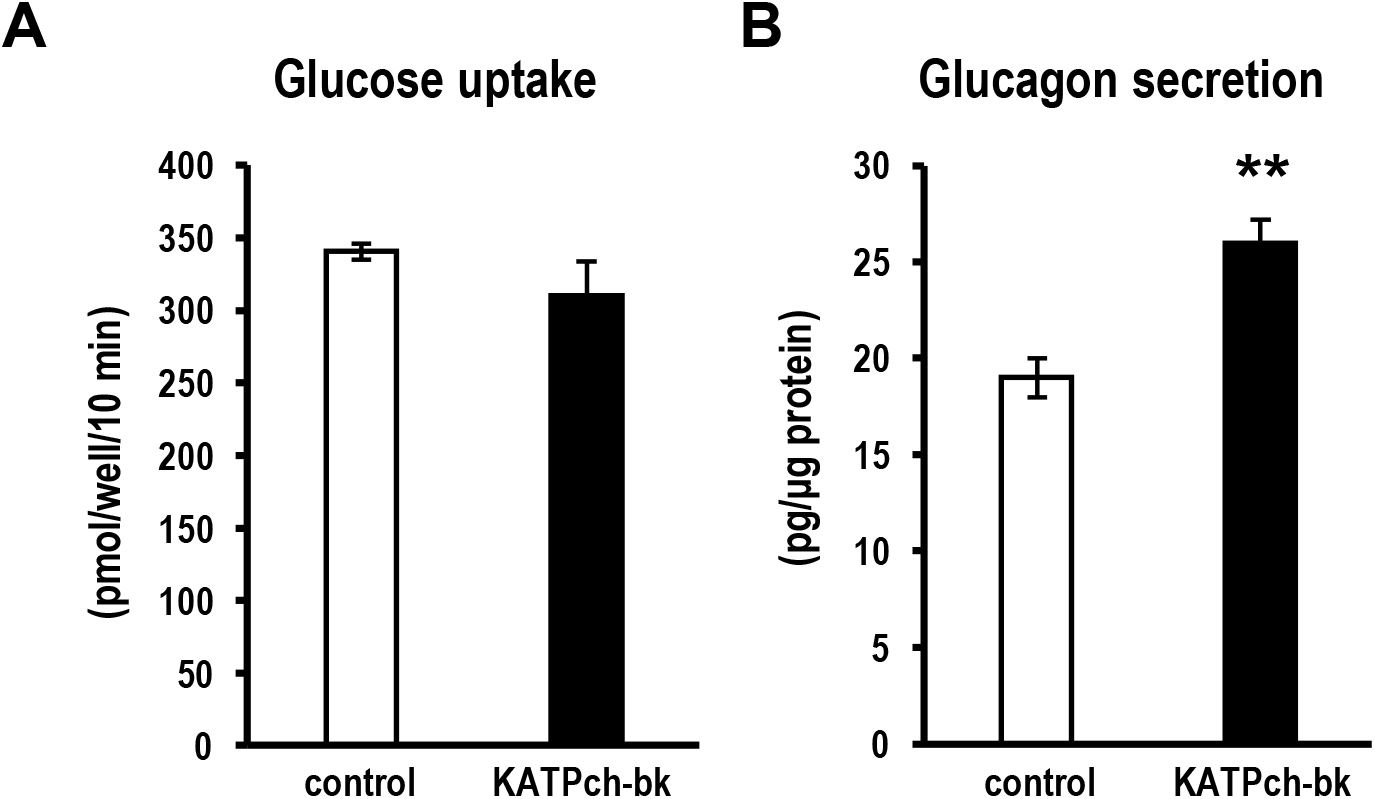
Pharmacological blockage of K/ATP channels increases glucagon secretion without affecting glucose uptake activity in alpha-TC1 cells. Uptake of 2-deoxy-D-glucose (A) and glucagon secreted in the cultured media (B), in the absence (open bar) or presence (closed bar) of an agonist of the sulfonylurea receptor on K/ATP channels. **; p < 0.01, vs control. n = 4-7.

### Intracellular ATP levels remain stable despite of the suppression of glucose uptake

Glucose serves as the predominant source for ATP production in the most of living cells. Given this pivotal role, we investigated how the suppression of glucose uptake impacted intracellular ATP levels upon exposure to the compounds. Neither the SGLT2 inhibitors dapagliflozin nor empagliflozin induced alterations in intracellular ATP levels. Similarly, cytochalasin B, which effectively suppressed glucose uptake, did not influence ATP levels (Fig. 5A). In addition, we confirmed that the K/ATP channel blockage had no discernible effect on ATP levels (Fig. 5B).

**Fig. 5.**
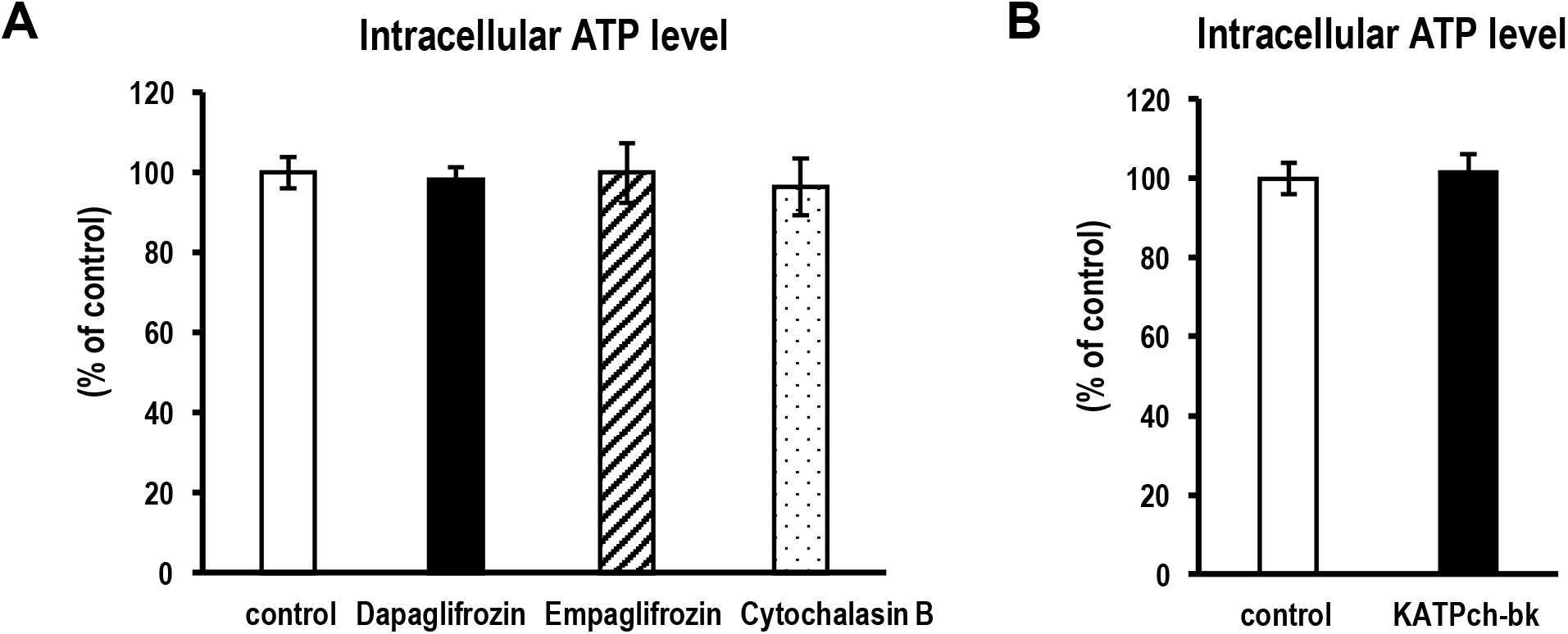
The suppressions of glucose uptake were not enough to reduce intracellular ATP pools. Amounts of intracellular ATP levels at the basal status (open bar) and after the inhibition of glucose uptake by dapagliflozin (closed bar), empagliflozin (hatched bar), or cytochalasin B (dotted bar) (A). The effects of K/ATP channel blockage on the intracellular ATP level are also shown (B). The vertical axis of the data represents the percentage of ATP levels relative to the control per well. No significant differences vs control. n=4.

## Discusstion

In recent years, there has been considerable debate regarding the presence of SGLT2 in pancreatic alpha cells as well as its possible role in the regulation of glucagon secretion. The direct role of SGLT2 in alpha-cell activity remains a point of contention in the literature. While some studies, such as those by Solini et al. and Kuhre et al., question the presence of SGLT2 in pancreatic alpha cells**[7,8]**, others have reported heterogeneous expression patterns, suggesting a more nuanced scenario**[5]**. These contrasting findings highlight the need for further investigation into the distribution and functional significance of SGLT2.

In this context, our study assessed the effects of SGLT2 inhibitors on glucose uptake and glucagon secretion in alpha-TC1 cells. We clearly demonstrated that dapagliflozin and empagliflozin, known as highly selective SGLT2 inhibitors, suppressed basal glucose uptake activity in alpha-TC1 cells even at moderate concentrations (Fig. 1). Our observations suggested that glucose transporters pharmacologically recognized as SGLT2 are present and functional in alpha-TC1 cells, supporting the notion proposed by Bonner et al. and Pedersen et al**[5,6]**. However, we failed to observe any significant alterations in glucagon secretion levels with these SGLT2 inhibitors, despite their direct action on alpha-TC1 cells (Fig. 2).

Pharmacological inhibition studies can often be confounded by compound specificity. However, the observed effects on glucose uptake and glucagon secretion can be attributed primarily to SGLT2 inhibition, given the high selectivity of the compounds used. Dapagliflozin and empagliflozin exhibit quite high selectivity for SGLT2 over SGLT1, with reported ratios of >1200 and >2500, respectively, in human cells **[16–18]**. Empagliflozin maintains high selectivity for mouse SGLTs, while it did not inhibit GLUT1 up to the concentration of 10 µM **[18]**. Even if SGLT1 is involved, our experimental results demonstrated that such highly selective SGLT2 inhibitors can significantly suppress glucose uptake in alpha-TC1 cells.

To further investigate this discrepancy, we successively examined to what extent the glucagon secretion would be affected by inhibition of SLC2A family glucose transporters by using a nonspecific inhibitor on GLUTs, cytochalasin B, as GLUT1 (SLC2A1), a primary member of the SLC2A, ubiquitously expresses and functions in almost all the mammal cells. In addition, the existence of GLUT1 in pancreatic alpha cells had been confirmed by a series of studies compared with SGLTs **[7–10]**.

Cytochalasin B inhibited basal glucose uptake by nearly 60%, confirming that GLUT-mediated glucose transport played a dominant role in alpha-TC1 cells. Despite this substantial inhibition, glucagon secretion showed only a slight decreasing trend without reaching statistical significance (Fig. 3). This finding contradicts previously proposed models in which changes in glucose uptake within alpha cells modulate glucagon secretion via effects on the K/ATP channel, suggesting instead that glucose uptake per se does not significantly influence glucagon secretion.**[19]** On the other hand, treatment with a weak K/ATP channel blocker markedly enhanced glucagon secretion without affecting glucose uptake (Fig. 4), thereby supporting the existence of the aforementioned model in which glucagon secretion is regulated via K/ATP channel activity.

To address this apparent contradiction, we measured intracellular ATP levels. We found that neither dapagliflozin nor empagliflozin altered ATP levels, and even cytochalasin B, which caused stronger inhibition of glucose uptake, had little to no effect on ATP levels (Fig. 5A, 3A). Mammalian cells rely heavily on ATP as the primary energy source for cellular activities, and intracellular ATP levels are maintained by robust homeostatic mechanisms. Our results indicated that, at least under acute conditions of glucose uptake inhibition, even a nearly 60% reduction in basal glucose uptake did not significantly alter intracellular ATP levels, and consequently did not affect K/ATP channel activity.

In summary, although SGLT2 appeared to function and be present in alpha-TC1 cells to a limited extent, its contribution to total glucose uptake was insufficient to alter intracellular ATP levels or glucagon secretion (Fig. 1, 5A). Therefore, we suggested that SGLT2-mediated glucose transport in alpha-TC1 cells does not significantly influence glucagon secretion (Fig. 2).

The expression of SGLT2 is often assessed by immunodetection using antibodies or by RT-PCR detection of mRNA. However, immunodetection with antibodies can be prone to artifacts, which is often a matter of debate **[7–11]**. Therefore, we performed reverse transcription using total RNA collected from alpha-TC1 cells and attempted to detect SGLT2 and SGLT1 expression by RT-PCR. Under standard PCR conditions (30 cycles), SGLT2 was below the detection limit, and SGLT1 was only barely detectable (data not shown). These findings are consistent with the view that SGLT2 is not expressed in alpha cells and that only SGLT1 is present. However, our observation that highly selective SGLT2 inhibitors significantly suppressed glucose uptake suggests that, at least in alpha-TC1 cells, SGLT2 is indeed expressed at a very low level but is nonetheless functionally active. Given the high selectivity of these inhibitors for SGLT2 over SGLT1, as well as the very low expression level of SGLT1 itself, it is reasonable to conclude that the effects of non-specific inhibition are minimal.

Although the alpha-TC1 cells used in this study may possess characteristics distinct from those of human pancreatic alpha cells—particularly with respect to SGLT function and the mechanisms regulating glucagon secretion—experimental studies using human islets face significant limitations. In particular, it is nearly impossible at present to study pure populations of human alpha cells in isolation. Increasing evidence indicates that the regulation of glucagon secretion involves not only intrinsic glucose-sensing mechanisms within alpha cells themselves, but also complex interactions with other endocrine cells in the islet, such as beta cells. Therefore, results obtained from experiments using purely isolated alpha cells should be interpreted separately from those using intact islets or in vivo models. Given that alpha-TC1 cells are, at present, the only widely available and well-established model of glucagon-secreting cells—and the only system in which pure populations of alpha cells can be maintained in culture—they occupy an indispensable position in glucagon research. Even if alpha-TC1 cells exhibit properties that differ from those of human alpha cells, elucidating these features remains crucial for advancing our understanding of alpha cell biology and glucagon regulation.

## Conclusion

In conclusion, our findings demonstrate that SGLT2 is functionally present at low levels in alpha-TC1 cells and can mediate basal glucose uptake; however, its pharmacological inhibition is insufficient to alter intracellular ATP levels or regulate glucagon secretion. These results support the notion that suppression of glucose uptake by highly selective SGLT2 inhibitors per se does not directly control glucagon secretion in this model, and highlight the predominant role of K/ATP channel activity in the regulation of glucagon release. Further studies using human islets and more physiologically relevant models are warranted to clarify the precise role of SGLT2 in alpha-cell physiology, especially in humans.

## Acknowledgments

We would like to express deep appreciation to Professors Tadahiro Kitamura (Gunma University) and Hiroyuki Mizuguchi (Osaka Ohtani University) for valuable advice related to glucagon measurement as well as culture of alpha-TC1 cells. We also thank Professor Hirokazu Miyoshi and Advance Radiation Research, Education, and Management Center of Tokushima University for advice and help for the experiments with radioisotopes, and Mrs. Koji Adachi, Chikashi Tokuda and Kazuki Sato for helpful advice related to glucagon assay based on HTRF technology.

## Graphical abstract

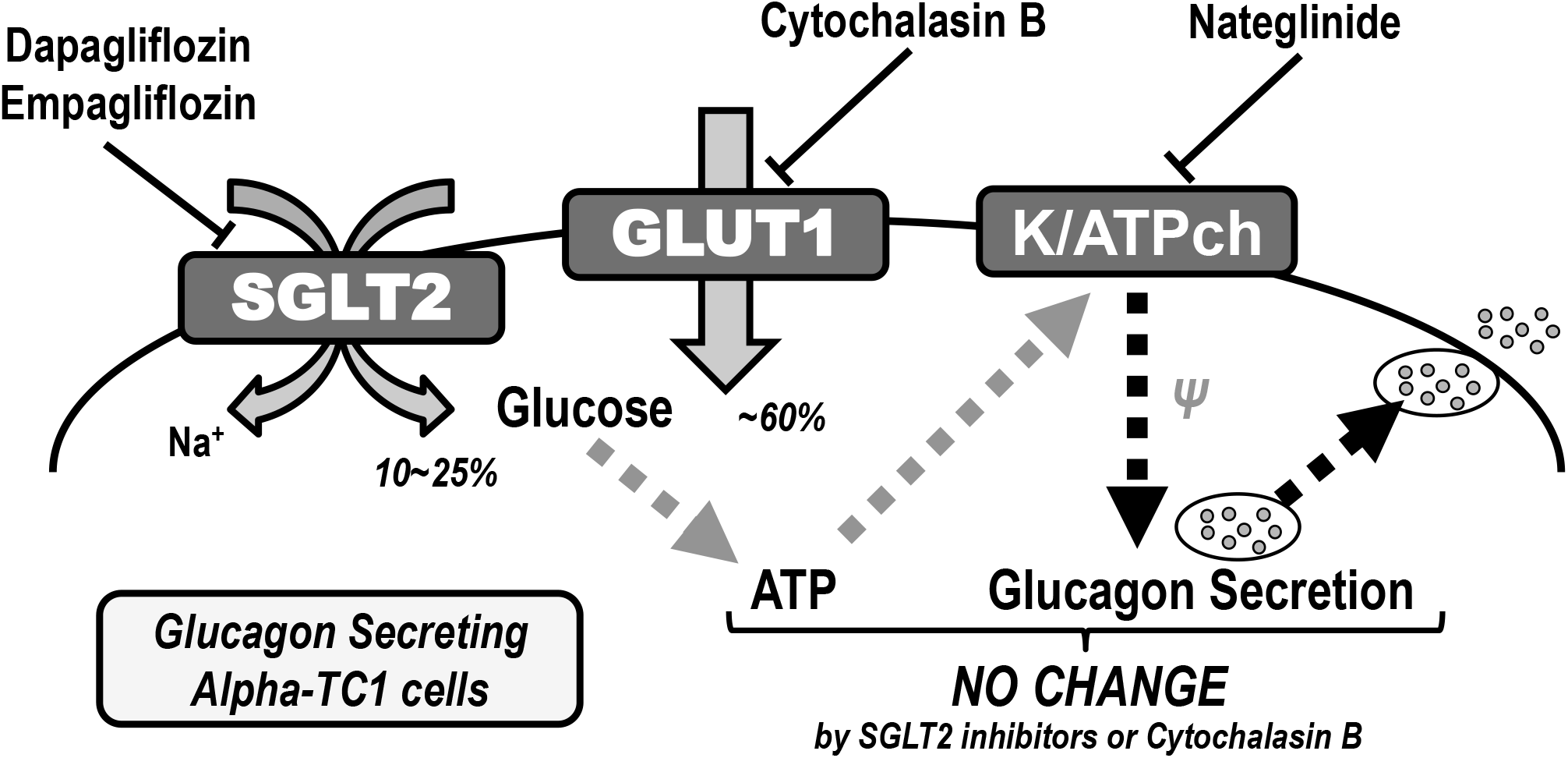

